# Cell-free Expression of Proton-Coupled Folate Transporter in the Presence of Nanodiscs

**DOI:** 10.1101/2021.05.19.444806

**Authors:** Hoa Quynh Do, Carla M. Bassil, Elizabeth I. Andersen, Michaela Jansen

## Abstract

The Proton-Coupled Folate Transporter (PCFT) is a transmembrane transport protein that controls the absorption of dietary folates in the small intestine. PCFT also mediates uptake of chemotherapeutically used antifolates into tumor cells. PCFT has been identified within lipid rafts observed in phospholipid bilayers of plasma membranes, a micro environment that is altered in tumor cells. The present study aimed at investigating the impact of different lipids within Lipid-protein nanodiscs (LPNs), discoidal lipid structures stabilized by membrane scaffold proteins, to yield soluble PCFT expression in an *E. coli* lysate-based cell-free transcription/translation system. In the absence of detergents or lipids, we observed PCFT quantitatively as precipitate in this system. We then explored the ability of LPNs to support solubilized PCFT expression when present during in-vitro translation. LPNs consisted of either dimyristoyl phosphatidylcholine (DMPC), palmitoyl-oleoyl phosphatidylcholine (POPC), or dimyristoyl phosphatidylglycerol (DMPG). While POPC did not lead to soluble PCFT expression, both DMPG and DMPC supported PCFT translation directly into LPNs, the latter in a concentration dependent manner. The results obtained through this study provide insights into the lipid preferences of PCFT. Membrane-embedded or solubilized PCFT will enable further studies with diverse biophysical approaches to enhance the understanding of the structure and molecular mechanism of folate transport through PCFT.

**Highlights:**

- Cell free expression of PCFT without any lipids or detergents resulted in quantitative precipitation of in-situ synthesized PCFT.
- Additives for expression of PCFT in the soluble fraction were identified.

## Introduction

Folate, also known as vitamin B9, is needed to synthesize purine, thymidylate, and methionine, which are essential in cell growth and division. In mammals, there is no *de novo* folate synthesis due to the absence of the folate synthase enzymes^1^. Instead, folate is obtained from dietary sources. Folate cannot permeate membranes directly due to its hydrophilic nature, and multiple membrane transport proteins, such as the folate receptor alpha (FRα)^2,3^, reduced folate carrier (RFC)^4^, and proton-coupled folate transporter (PCFT)^5–7^, transport this molecule into mammalian cells. PCFT is the primary transporter for dietary folate uptake in the gut and has a high expression level in the proximal small intestine^5,8^, i.e., in the duodenum^9^ and the jejunum^10^. It is also found to be expressed in epithelial cells of the choroid plexus, where it aids the FRα-mediated transport of folate into the epithelial cytosol^11,12^. Folate is then further transported to the cerebrospinal fluid by RFC. Loss-of-function mutations in PCFT, the mediator of the intestinal absorption and delivery of folate to the central nervous system^7,13,14^, cause hereditary folate malabsorption^13,15–20^ with clinical and biochemical features such as diarrhea, anemia, leukopenia, cognitive and motor impairment, pneumonia, and undetectable folate levels in CSF^15,21,22^. Cerebral folate deficiency is also connected to in Alzheimer’s disease^23–27^ and inferred to cause epilepsy, autism spectrum disorders, and other neurological disorders in young children^28,29^. Treatment of cerebral folate deficiency with folinic acid^20,28,30,31^ and its S-enantiomer levofolinic acid^22^ results in clinical improvement in most patients who receive the treatment before the age of six. Folinic acid is a reduced folic acid derivative, the 5-formyl tetrahydrofolic acid, with complete vitamin activity. However, for some patients with early treatment and others with a treatment delayed beyond six years of age, incomplete neurological recovery or continuous neurological deficits have been reported^22,28^. Therefore, toward the future development of more effective therapies for neurological disorders involving PCFT mutations, it is essential to understand the atomic structure of PCFT, which is currently unavailable.

PCFT functions optimally at an acidic milieu^5,8,32^, approximating the microenvironment of the proximal intestine and also solid tumors. Recent findings reveal the upregulation of PCFT with the highest levels in the cells derived from colorectal adenocarcinoma, ovarian carcinoma, hepatoma, and small cell lung cancer cell lines^33^. Unlike RFC, PCFT exhibits high affinity for both folic acid and 5-methyltetrahydrofolate at acidic pH (K_m_ ~ 0.5~1 μM), and it is thus likely that PCFT is the main route by which folate enters into cancer cells. These distinctions render PCFT as an ideal candidate for targeting solid tumors.

PCFT is a member of the Major Facilitator Superfamily and contains 12 α-helical transmembrane segments^10,34,35^. To facilitate future structure-function studies of PCFT that require pure, soluble, and functional PCFT, this study explores the impact of different lipids within preformed lipid-protein nanodiscs (LPNs) on the soluble, cell-free expression of PCFT into these LPNs.

## Materials and Methods

### Reagents

The pEXP5-NT/TOPO vector (Thermo Fisher) and the S30 T7 High-Yield Protein Expression System (Promega) were used to express proton-coupled folate transporter (PCFT) in vitro. POPC (Avanti Polar Lipids), DDM (Anatrace), Tween 20 (Fisher Scientific), Triton X-100 (Sigma-Aldrich), and lipid-protein nanodiscs (MSP1D1-His-POPC, MSP1D1-His-DMPG, MSP1D1-His-DMPC, Cube Biotech) were used to solubilize in-situ synthesized PCFT. A rabbit polyclonal 6X-His tag antibody conjugated to horseradish peroxidase (ab1187, Abcam) was used to detect PCFT and MSP1D1.

### Preparation of plasmid DNA

The open reading frame for full-length human proton-couple folate transporter (PCFT), UniProtKB accession number Q96NT5, was cloned into the pEXP5-NT/TOPO vector to obtain PCFT-pEXP5-NT. The DNA sequence of the resulting PCFT construct contains an N-terminal 6X-His tag followed by a Tobacco Etch Virus (TEV) protease recognition site. The sequence was confirmed by DNA sequencing (Genewiz). Plasmid DNA was isolated with the EndoFree Plasmid Maxi Kit (Qiagen) following the manufacturer’s protocol.

### Cell-free expression of PCFT - Protein Synthesis Reaction

The S30 T7 High-Yield Protein Expression System was used for cell-free PCFT protein expression in the presence and the absence of LPNs, lipids, and detergents. This expression system is an *Escherichia coli* extract-based cell-free protein synthesis system, which consists of T7 RNA polymerase for transcription and all necessary components for translation.

PCFT-pEXP5-NT was used as template, which contains a T7 promoter, a ribosome binding site, and the DNA sequence encoding for full-length human PCFT protein (Fig. 1). The typical reaction volume was 5 μL using the template, T7 S30 Extract for circular templates, and S30 Premix Plus as recommended by the manufacturer (Fig. 2). Reactions were incubated in a thermomixer at 1,200 rpm and 37°C for 1 hour.

**Figure 1.**
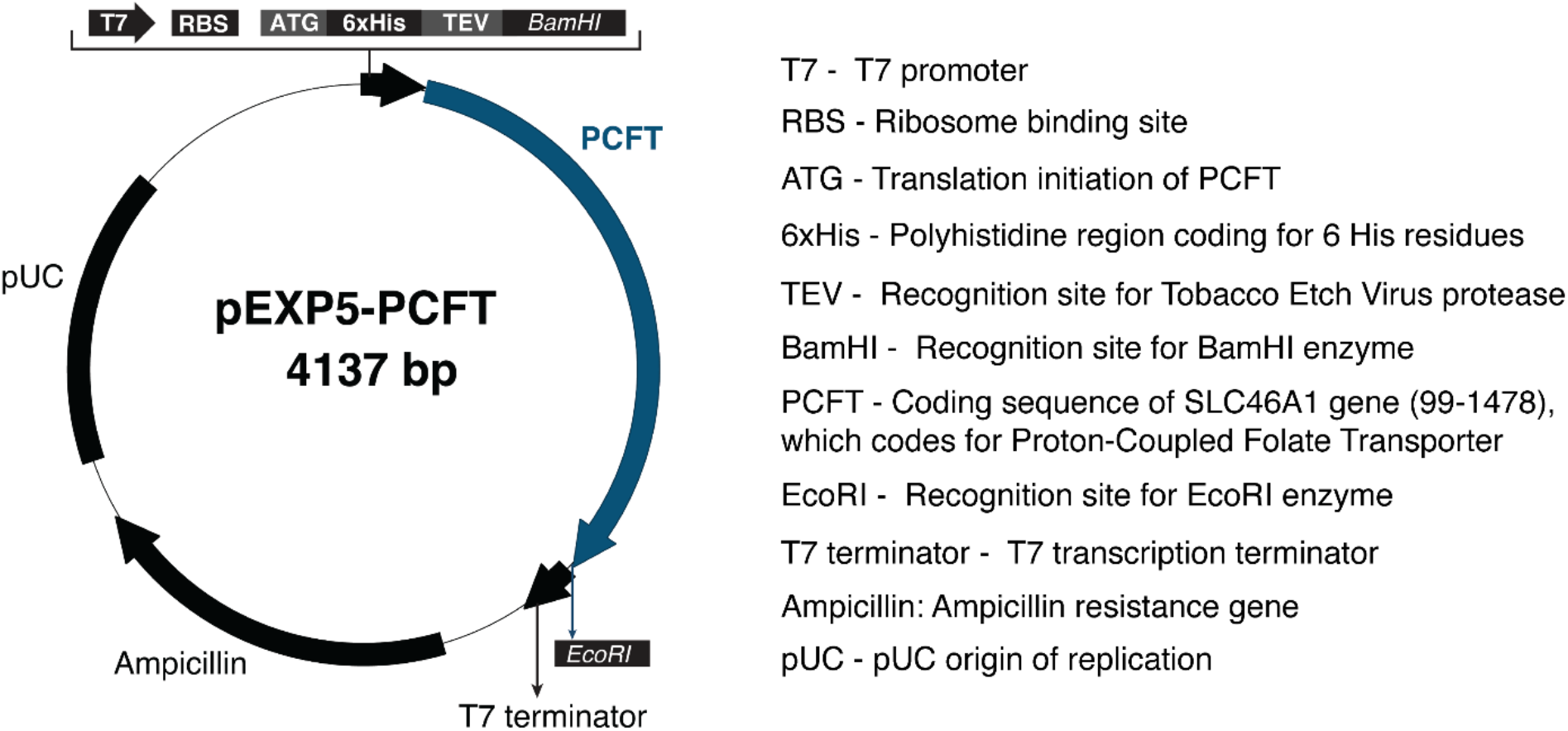
PCFT expression plasmid. The coding sequence of SLC46A1 gene, nucleotides from 99 to 1478, together with two recognition sites for BamHI and EcoRI restriction enzymes, were cloned into pEXP5-NT/TOPO vector purchased from ThermoFisher Scientific. The correct DNA sequence of the PCFT construct was confirmed by Genewiz.

**Figure 2.**
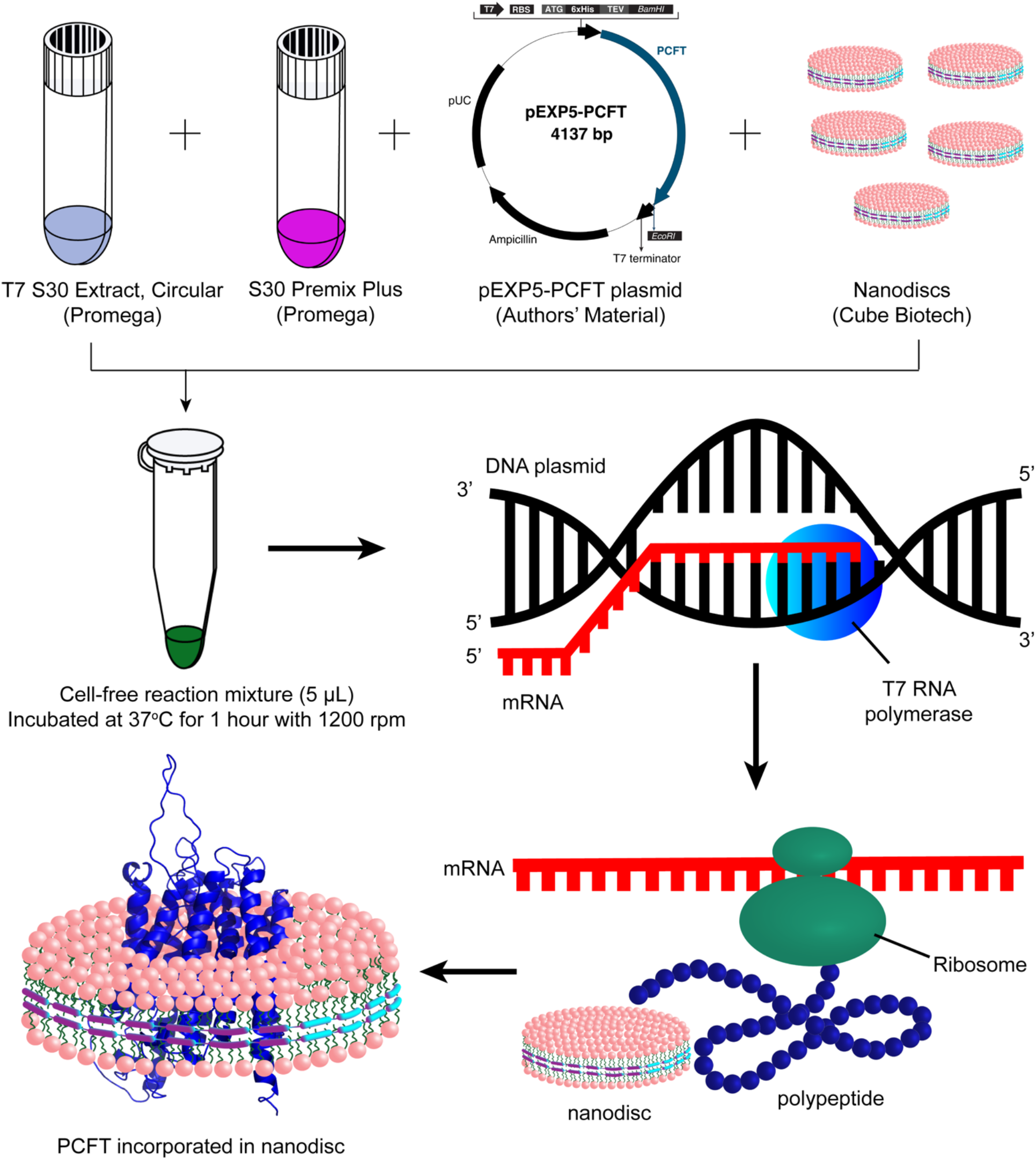
Schematic for Cell-free expression of PCFT in the presence of nanodiscs. T7 S30 Extract, Circular, and S30 Premix Plus are components of the S30 T7 High-Yield Protein Expression System purchased from Promega. The pEXP5-PCFT plasmid is the PCFT construct shown in Fig. 1. Empty LPNs containing POPC, DMPG, or DMPC were added to a cell-free reaction mixture. Soluble PCFT was observed when PCFT was cell-free expressed in the presence of the nanodisc.

Pre-assembled LPNs and/or detergents were added to some reactions. PCFT was expressed in the presence of pre-assembled LPNs consisting of the membrane scaffold protein MSP1D1-His with a single phospholipid; 1-palmitoyl-2-oleoyl-sn-glycero-3-phosphocholine (POPC), 1,2-dimyristoyl-sn-glycero-3 - phosphocholine (DMPG), or 1,2-dimyristoyl-sn-glycero-3-phosphocholine (DMPC), MSP1D1-His-POPC, MSP1D1-His-DMPG, and MSP1D1-His-DMPC, respectively. Three different LPN concentrations were used following the LPN manufacturer’s manual, 20 μM, 40 μM, and 80 μM. The mixtures were incubated for 1 hour at 37°C before the separation of soluble and insoluble PCFT.

After the expression, each sample was centrifuged at 22,000 g for 10 min at 4°C to separate insoluble PCFT in the pellet from soluble PCFT in the supernatant. Each pellet fraction was resuspended in 50 μL of 1x Laemmli sample buffer containing 1% SDS, with or without 2.5 % of 2-mercaptoethanol. Each supernatant fraction was diluted using the same sample buffer in a total volume of 50 μL.

### Immunoblot analysis

Samples from the pellet and supernatant fractions were heated at 70°C for 10 minutes to enhance protein denaturation. Ten microliters of each sample, which corresponds to 1 μL of the initial reaction volume, were separated on Mini-PROTEAN TGX gels (BioRad) and transferred to PVDF membranes (BioRad) using the Bio-Rad trans-blot turbo system following the manufacturer’s protocol. PVDF membranes were briefly immersed in methanol and then equilibrated in western transfer buffer (20% of Bio-Rad 5x transfer buffer, 20% ethanol) for 3-5 minutes. Proteins were transferred at 2.5 Amperes and up 25 Volts for 3 minutes at room temperature. Polyvinylidene fluoride (PVDF, Biorad) membranes were blocked under agitation in 5% nonfat milk in tween-containing tris-buffered saline buffer (TTBS buffer: 0.1% Tween-20, 100 mM Tris, 0.9% NaCl, pH 7.5) for one hour or overnight. Afterwards, the blot was incubated with 6X-His tag antibody conjugated with HRP in a dilution of 1:5000 for one hour under gentle agitation. Subsequently, the blot was washed five times, each for 5 minutes with 5 ml of TTBS buffer. After an additional wash with tris-buffered saline (100 mM Tris, 0.9% NaCl, pH 7.5), SuperSignal West Femto Maximum Sensitivity Substrate (Thermo Scientific) was used for imaging with a digital imaging system (ImageQuant™ LAS 4000, GE Healthcare).

### Data Analysis

PCFT quantification was performed to determine if PCFT solubility depends on the lipid composition and concentrations of LPNs. The soluble PCFT bands on western blot images were selected, the band intensities were plotted as peaks, and the area under a peak was analyzed using ImageJ software. The resulting values were normalized to the highest value obtained from soluble PCFT expressed in LPNs. Statistical significance in PCFT solubility, depending on lipid concentrations and compositions, was determined using one-way ANOVA with Tukey’s multiple-comparisons test in Prism 6 Software (GraphPad Prism, La Jolla, CA).

Data charts were generated using Microsoft Excel and Prism 6 Software (GraphPad Prism, La Jolla, CA). Figures were created using ApE plasmid editor, Adobe Illustrator 2020, and Adobe Photoshop 2020.

## Results

A detailed molecular understanding of proton-coupled folate transporter (PCFT) is desirable and essential in the structure-based design of drugs and therapies effectively targeting folate-related neurological disorders and cancer. Biochemical and biophysical techniques routinely applied to study the structure and function of proteins frequently require soluble or solubilized (membrane) proteins. For integral membrane proteins a critical step in making protein available for biophysical approaches such as for example cryoelectron microscopy, circular dichroism, isothermal titration calorimetry, or X-ray crystallography is the solubilization of the protein from the native lipid bilayer or the integration into a stable bilayer (mimetic) or detergent micelle^36^. To produce soluble PCFT, a PCFT construct was designed as part of this study, containing full-length human PCFT cloned into the pEXP5-NT/TOPO vector (Fig. 1). The cell-free transcription and translation system was then used to examine the expression and solubility of PCFT in the presence (Fig. 2) or the absence of LPNs containing different lipids.

### PCFT expression is observed in cell-free expression system

Transcription/translation reactions contained PCFT-pEXP5-NT as a template. After incubation reactions were separated by centrifugation into pellet and supernatant fractions. Since PCFT is a transmembrane protein, in the absence of detergent or an otherwise solubilizing milieu, the translated protein is expected to be in the pellet, whereas soluble PCFT would be in the supernatant fraction (Fig. 3). Proteins in both fractions were separated on SDS-PAGE and detected using western blotting with an antibody directed against the N-terminal 6X-His tag present in the PCFT construct.

**Figure 3.**
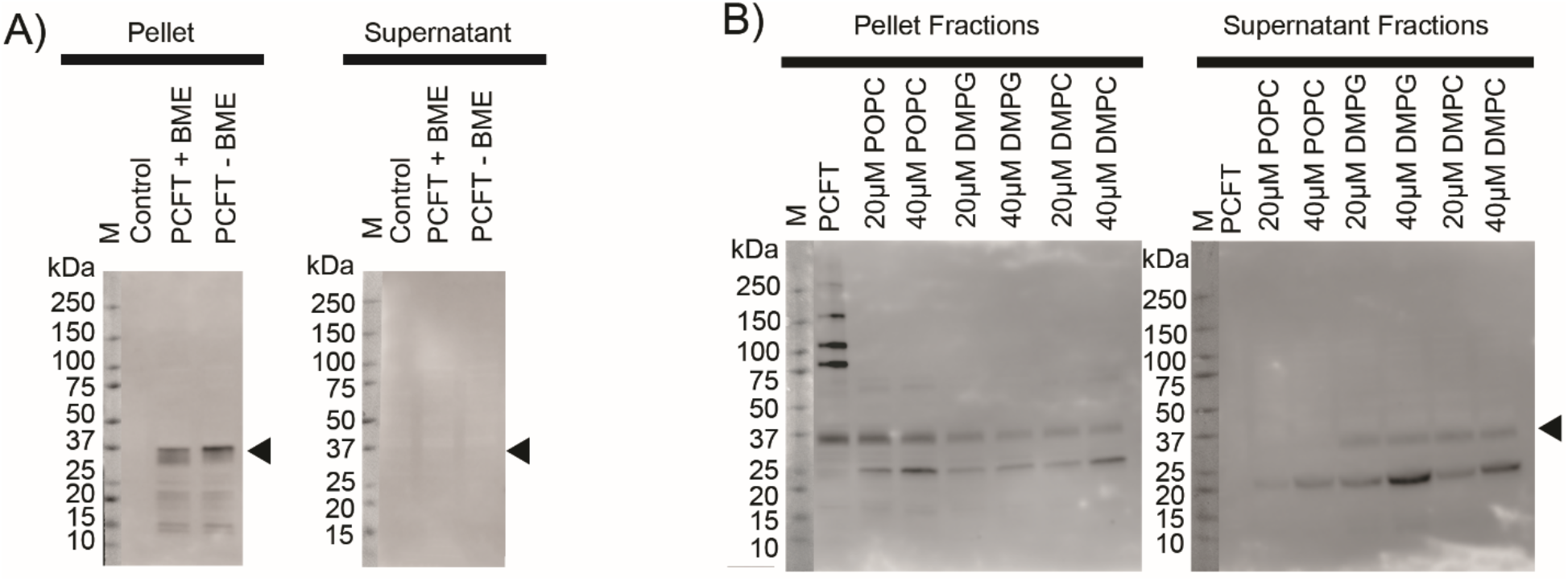
Cell-free expression of proton-coupled folate transporter (PCFT). The arrows indicate the position of monomeric PCFT (~37 kDa). A) Cell-free expression of PCFT in the absence of LPNs. The “Control” lane reactions did not contain PCFT-pEXP5-NT plasmid. *β-mercaptoethanol* (BME) was added to some reactions as indicated. B) Cell-free expression of PCFT in the presence of 20 μM or 40 μM LPNs containing POPC, DMPG, or DMPC lipids. The “PCFT” lane reactions did not contain any LPNs.

In the absence of the plasmid PCFT-pEXP5-NT template, no band, and consequently no PCFT protein, was detected in either the pellet or supernatant fractions (Fig. 3A, lanes ‘Control’), whereas the presence of PCFT-pEXP5-NT template in the reaction yielded a band at approximately 37 kDa (Fig. 3A, pellet fractions). The 6X-His-tagged full-length human PCFT construct in this work consists of 493 amino acids with a theoretical molecular weight of 52 kDa. When expressed in *Spodoptera frugiperda* (Sf9) insect cells^37^, Chinese hamster ovary cells^38^, or *Xenopus laevis* oocytes^38^ monomeric PCFT migrates in two bands between 35 and 55 kDa. The electrophoretic mobility of PCFT depends on the post-translational modification of N-linked glycosylation^39^. Removing the glycosylation with PNGase F results in PCFT that mainly migrates at ~35 kDa^39^. The migration of PCFT observed here (~37 kDa) is consistent with nonglycosylated PCFT as expected in the cell-free expression system where post-translational modifications do not occur.

This data indicates that the PCFT protein was expressed in the cell-free expression system using PCFT-pEXP5-NT as a template. As would be expected in the absence of detergents or bilayers, the expressed PCFT is exclusively insoluble, and PCFT protein is only found in the pellet but not the supernatant fraction.

### Cell-free expressed and soluble PCFT is detected in the presence of LPNs

Next, the cell-free expression of PCFT was examined in the presence of LPNs of differing lipid compositions. LPNs were stabilized by the membrane scaffold protein MSP1D1 (genetically modified apolipoprotein A1 with N-terminal globular region replaced by 6X-His and Tobacco Etch Virus protease site). The ~10-nm size of MSP1D1 LPNs was used to be tailored to the diameter of a transmembrane protein with 12 α-helical transmembrane segments. Fig. 3B shows PCFT translation using cell-free expression in the presence of MSP1D1 LPNs containing zwitterionic POPC with a saturated palmitoyl- and unsaturated oleoyl chain (16:0-18:1 PC; 1-palmitoyl-2-oleoyl-sn-glycero-3-phosphocholine), zwitterionic saturated DMPC (14:0 PC; 1,2-dimyristoyl-sn-glycero-3-phosphocholine) or anionic saturated DMPG (14:0 PG; 1,2-dimyristoyl-sn-glycero-3-phospho-(1’-rac-glycerol)) (Cube Biotech) with PCFT-pEXP5-NT (6X-His-PCFT) as a DNA template. In the absence of LPNs, once again only insoluble PCFT was observed. In the presence of 20 or 40 μM LPNs, PCFT was found in both insoluble form in the pellet and soluble form in the supernatant (Fig. 3B). More soluble PCFT was detected for reactions with LPNs containing the saturated lipids DMPG or DMPC as compared to those containing unsaturated POPC.

### DMPG and DMPC LPNs produce soluble PCFT

To further assess the impact of LPNs, their concentrations, and lipid compositions on the PCFT solubility, the experiment in Fig. 3B was repeated with higher concentrations of LPNs, i.e., 80 μM LPNs, containing POPC, DMPG, and DMPC lipids (Fig. 4A). ‘PCFT’ lanes represent PCFT expressed in the absence of LPNs, and the lack of soluble PCFT again confirms that without LPNs, only insoluble PCFT was obtained in the cell-free expression. When LPNs were added to the reaction after transcription and translation, no soluble PCFT was found (data not shown). In contrast, depending on the specific lipid within LPNs, both insoluble and soluble PCFT were observed when LPNs were present during transcription and translation in the expression reaction. No or minimal PCFT was incorporated into POPC LPNs. For all three POPC LPN concentrations used, 20, 40, and 80 μM, when the band intensity was assessed using one-way ANOVA with Tukey’s multiple-comparisons test, the PCFT band intensities were not significantly different from PCFT expressed in the absence of any solubilizing agent. At a concentration of 20 μM, DMPG LPNs yielded approximately three times the soluble PCFT intensity as compared to POPC LPNs. The soluble PCFT in DMPG reactions was significantly above the reactions without LPNs (p ≤ 0.001, p ≤ 0.01, p ≤ 0.01 for 20, 40, and 80 μM, respectively). However, the concentration of the DMPG LPNs itself did not impact the soluble PCFT in a concentration-dependent manner.

**Figure 4.**
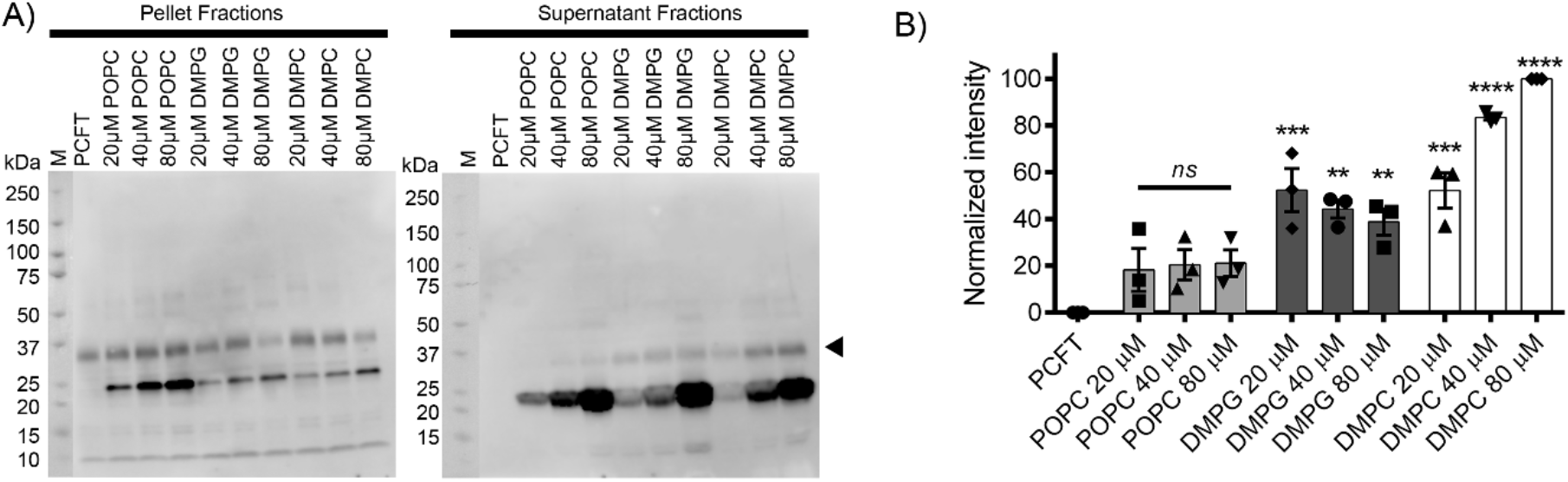
Solubilization of PCFT in different lipid concentrations and compositions. A) Cell-free expression of PCFT in the presence of 20 μM, 40 μM, or 80 μM LPNs containing POPC, DMPG, or DMPC lipid. PCFT lane did not contain any LPNs. B) PCFT solubilization in 20 μM, 40 μM, or 80 μM LPNs containing POPC, DMPG, or DMPC lipid. The PCFT band intensity was quantified in western blots of supernatant fractions and normalized to the PCFT band derived from 80 μM DMPC LPNs. The data show the mean ± SEM from n=3 independent experiments. The highest level of solubilized PCFT was found in LPNs containing DMPC lipid. LPNs containing POPC lipid yielded minimal levels of solubilized PCFT. Statistical significance was determined using was determined using one-way ANOVA with Tukey’s multiple-comparisons test in Prism 6 Software (GraphPad Prism). Significance is indicated vs “PCFT” samples without LPNs as ** p ≤ 0.01, *** p ≤ 0.001, and p ≤ ****0.0001). Nonsignificant p value is shown as ns.

Similar to 20 μM DMPG LPNs, the same concentration of 20 μM DMPC yielded a three-fold increase in soluble PCFT as compared to POPC LPNs. Contrary to DMPG LPNs, DMPC LPNs led to a concentrationdependent increase in soluble PCFT. The highest concentration of DMPC LPNs of 80 μM, produced the highest amount of soluble PCFT, approximately two-fold higher than all DMPC concentrations.

Overall, incorporation of PCFT into LPNs containing DMPG and DMPC (the more ordered or condensed lipids compared to POPC^40^) was higher than compared to POPC LPNs with an overall increase in solubility from POPC < DMPG < DMPC. PCFT has been found in lipids rafts. Reduced levels in lipid rafts have been associated with folate malabsorption in chronic alcoholism^41^. Changes to the lipid raft composition in cancer have been implicated in regulating cell proliferation, apoptosis, and cell migration^42^. Lipid rafts are microdomains found in the plasma membrane that are enriched in sphingolipids and cholesterol. High levels of cholesterol in membranes lead to directional organization of the lipid bilayer due to the rigid sterol^43^. The packing of cholesterol in between lipid acyl chains consequently results in a closer packing or compaction of the lipid bilayer, a state of the bilayer also termed as “liquid-ordered”. Therefore, the preference of PCFT to integrate into LPNs containing ordered lipids is comparable to its native localization within lipid raft microdomains.

## Discussion and Conclusion

PCFT was identified as the main pathway for dietary folate uptake in the proximal small intestine in 2006^13^. Detailed studies of PCFT using PCFT constructs modified with amino acids engineered at specific positions, including the use of the substituted cysteine accessibly method (SCAM)^44^, have provided insights into the topology and conformational transitions during substrate translocation^16,18,35,38,45–54^. However, much remains unknown about PCFT’s detailed molecular structure and substrate translocation mechanism.

In this study PCFT was produced in a soluble form using cell-free synthesis in the presence of membrane-mimicking LPNs. Soluble PCFT was obtained with a continuous one-step reaction and within one hour. In preliminary experiments, detergent micelles alone or mixed detergent-lipid micelles did not support soluble PCFT synthesis. The described soluble PCFT expression directly into LPNs facilitates the expression of engineered constructs with site-specific modifications. The expression level is amenable for diverse biophysical approaches to study PCFT’s structure and function, such as applications using fluorescence (tryptophan fluorescence or single-molecule fluorescence resonance energy transfer, smFRET), cryo-EM, isothermal titration calorimetry or circular dichroism.

## Abbreviations

(PCFT): Proton-coupled folate transporter
(FRα): folate receptor alpha
(RFC): reduced folate carrier
(HRP): horseradish peroxidase
(MSP): membrane scaffold proteins
(DDM): N-dodecyl β-d-maltoside
(POPC): 1-palmitoyl-2-oleoyl-sn-glycero-3-phosphocholine
(DMPG): 1,2-dimyristoyl-sn-glycero-3-phosphocholine
(DMPC): 1,2-dimyristoyl-sn-glycero-3-phosphocholine
(BME): 2-mercaptoethanol

## Author Contributions

M.J. designed research. C.B and E.A. performed experiments. H.D. and M.J. analyzed data. H.D. and M.J. wrote the article with the input of all other authors.

## Acknowledgments

We thank the TTUHSC Core Facilities; some of the images and/or data were generated in the Image Analysis Core Facility and Molecular Biology Core Facility supported by TTUHSC. Research reported in this publication was supported in part by the TTUHSC Office of Research, and the Laura W. Bush Institute for Women’s Health & UMC Health System with seed grants, and by the National Institute of Neurological Disorders and Stroke of the National Institutes of Health under award number R01/R56NS077114 (to M.J.).

## Notes

### Competing Interest Statement

The authors have declared no competing interest.

